# Next-Gen Profiling of Tumor-resident Stem Cells using Machine Learning

**DOI:** 10.1101/2023.11.24.568600

**Authors:** Debojyoti Chowdhury, Bhavesh Neekhra, Shreyansh Priyadarshi, Swapnanil Mukherjee, Debashruti Maity, Debayan Gupta, Shubhasis Haldar

## Abstract

Tumor-resident stem cells, also known as cancer stem cells (CSCs), constitute a subgroup within tumors, play a crucial role in fostering resistance to treatment and the recurrence of tumors, and pose significant challenges for conventional therapeutic methods. Existing approaches for identifying CSCs face notable hurdles related to scalability, reproducibility, and technical consistency across different cancer types due to the adaptable nature of CSCs. In this context, we introduce OSCORP, an innovative machine-learning-driven approach. This methodology quantifies and identifies CSCs, achieving almost 99% accuracy using biopsy bulk RNAseq data. OSCORP leverages genetic similarities between normal and cancer stem cells. By categorizing CSCs into four distinct yet dynamic potency states, this approach provides insights into the differentiation landscape of CSCs, unveiling previously undisclosed facets of tumor heterogeneity. In evaluations conducted on patient samples across 22 cancer types, OSCORP revealed clinical, transcriptomic, and immunological signatures associated with each CSC state. It has emerged as a comprehensive tool for understanding and addressing the complexities of cancer stem cells. Ultimately, OSCORP opens up new possibilities for more effective personalized cancer therapies and holds the potential to serve as a clinical tool for monitoring patient-specific CSC changes during treatment or follow-up care.

## Introduction

Tumor-resident stem cells (or cancer stem cells, CSCs) constitute a distinctive subset within tumors, mirroring normal stem cells in their extraordinary capacity for self-renewal and differentiation [1,2]. The adaptive capabilities of CSCs under selective pressures give rise to a plethora of diverse subpopulations, significantly contributing to the observed tumor heterogeneity [7,16]. This inherent diversity becomes a driving force for the evolution of CSCs through a dynamic interplay of random mutations and epigenetic alterations, facilitating their adept navigation within the dynamic tumor microenvironment [17,18]. Embracing the principles of natural selection, the plasticity of CSCs serves as a critical determinant in shaping tumor progression and confounding treatment strategies, thereby promoting evasion [3,19]. The survival advantage of CSCs, proficient in resource utilization and adaptation to posttreatment conditions, ultimately manifests in tumor recurrence at either the primary or newly developed sites [20–22]. Additionally, CSCs exhibit a phenotypic signature characterized by heightened migratory and invasive properties, actively contributing to the metastatic cascade and underscoring their pivotal role in cancer progression [4,19]. The imperative for real-time, precise quantification and identification of CSCs during treatment becomes evident, offering profound insights into their dynamic behavior and enabling the strategic tailoring of treatment approaches to counter resistance and mitigate the risk of relapse. A comprehensive exploration of the quantitative landscape of CSCs has the potential to unveil their intrinsic diversity, significantly advancing our nuanced understanding of cancer biology [23]. Pioneering the identification of CSCs based on their differentiation potential opens new avenues for the exploration of innovative therapeutic interventions, directing focus toward specific CSC states and enhancing treatment efficacy for improved patient outcomes [5,24]. This multifaceted perspective on CSCs underscores the intricate complexities of cancer progression and treatment resistance, providing a foundation for tailored interventions in the pursuit of improved clinical outcomes.

Current investigations often grapple with a comprehensive understanding of tumor-resident stem cells [8,9]. Despite advancements in addressing tumor stemness [25], uncertainties persist in accurately identifying CSCs, deciphering their intricate interplay with clinical features, and uncovering their genomic diversity across diverse cancer types [8]. Further elucidation is essential regarding the mechanisms governing the transition of CSCs between distinct states and their dynamic interactions within the tumor ecosystem [26,27]. The intrinsic dynamism, phenotypic adaptability, and heterogeneity of CSCs pose significant challenges for existing quantification methods [8–10]. While marker-based approaches contribute valuable insights, they fall short of capturing the complete spectrum of CSC characteristics [11,28]. This underscores the necessity for pioneering strategies that integrate both functional and phenotypic dimensions of CSCs, ensuring precise identification and the potential for targeted therapeutic interventions. The intricacies of CSC biology demand ongoing exploration, emphasizing the need to refine methodologies, thereby enhancing our understanding and paving the way for more effective treatment strategies in the field of cancer research.

At the forefront of cutting-edge cancer research, artificial intelligence (AI) stands out as a powerful tool, providing a robust means to unravel the intricacies embedded within CSC genomic data [25,29–31]. AI algorithms demonstrate unparalleled proficiency in navigating intricate datasets, unveiling subtle patterns, and discerning rare cellular subpopulations often overlooked by conventional methods [32,33]. Utilizing the paradigm of computational cytometry [33], our innovative approach, Optimized digital cancer Stem Cell sORter and Profiler (OSCORP), leverages the power of machine learning to traverse extensive patient datasets. It effectively quantifies elusive CSC subpopulations and identifies nuanced patterns with an exceptional accuracy rate of 98.87%. Introducing a novel CSC classification into four genetically distinct yet dynamically entropic states that mirror various human stem cell types (iPSCs, hESCs, hMSCs, and hUSCs), OSCORP enriches our understanding of their roles in cancer progression. Tumor plasticity, assessed through Markov chain entropy (MCE), sheds light on CSC differentiation and dedifferentiation potential, resembling a Waddington-like landscape. When applied comprehensively across 22 TCGA cancer types, these insights reveal intricate genomic, clinical, and immunological signatures intricately linked to diverse CSC populations. By unraveling the heterogeneity of CSCs and their profound implications, our findings propose a scientific classification of CSCs into four distinctive states based on their potency. This presents an opportunity to tailor therapeutic strategies toward personalized interventions and enhance prognostication.

## Results

### OSCORP can quantify CSCs and identify their potency state

OSCORP was intricately developed to achieve precise deconvolution of bulk RNAseq data, with the primary objective of discerning various cell types within tumors, notably CSCs (Fig. 1a). To accomplish this, we meticulously crafted a signature matrix encompassing a diverse array of leukocyte subsets, 22 distinct types of malignant cells, and CSCs (encompassing pluripotent, multipotent, and unipotent stem cell data) (Extended Data Fig. 1a). This singular cell-derived signature matrix was then leveraged for deconvolving RNA-seq data across 22 cancer types within The Cancer Genome Atlas (TCGA, n= 8786). Operating on an algorithm initially established by Newman *et al*. and validated using known cell populations from flow cytometric analysis33, our platform employs a support vector regression-based quantifier module. This module not only serves as a real-time CSC quantification tool but also unravels tumor heterogeneity by exposing the frequencies of diverse subpopulations. This sophisticated approach provides nuanced insights into the distinctive roles and contributions of various cell populations, CSCs included within each tumor sample, enriching our understanding of the intricate cellular dynamics at play.

**Figure 1.**
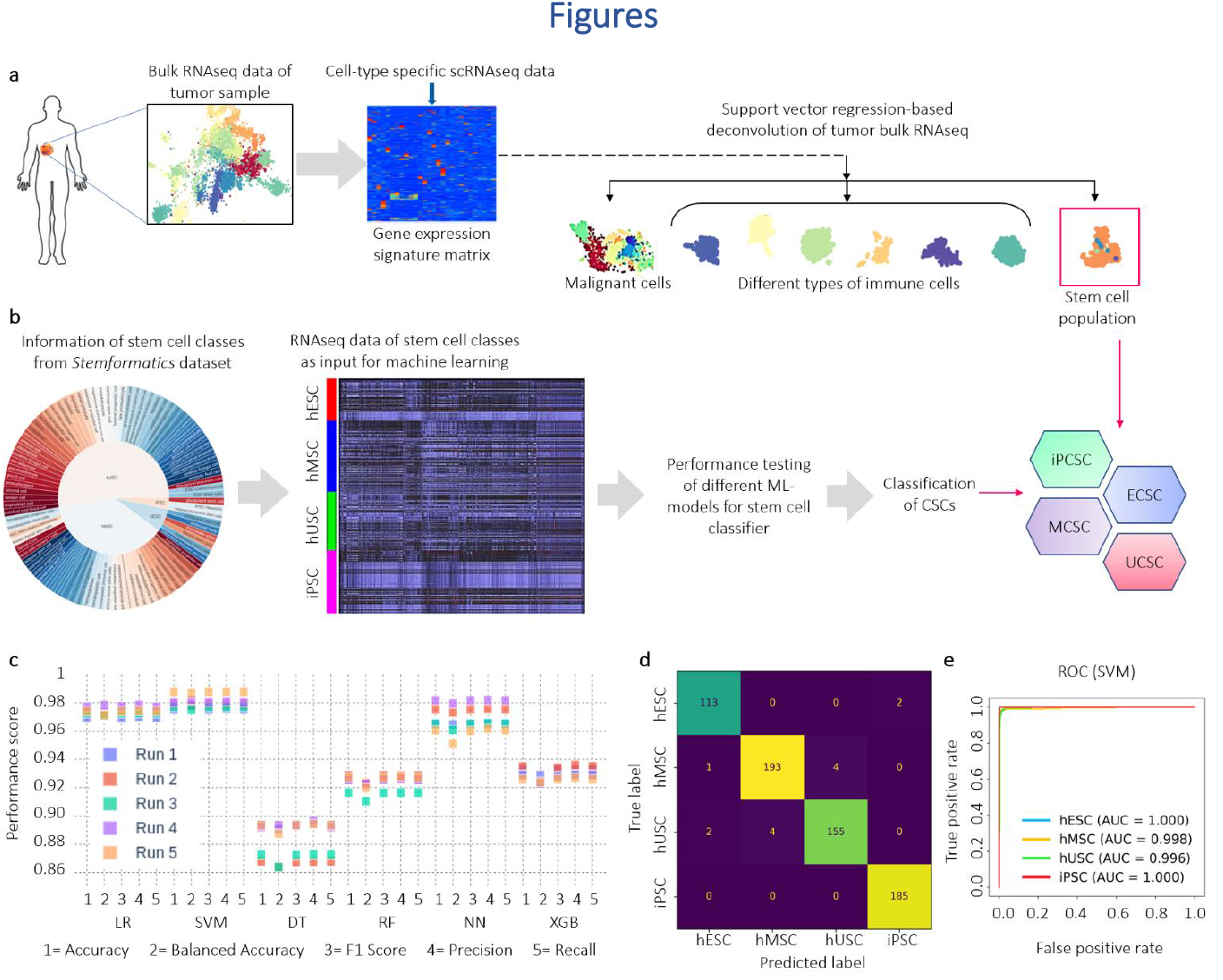
ML-based approach to quantify and identify cancer stem cells from tumor sample genomic data. a. Workflow of deconvoluting tumor samples utilizing support vector regression (SVR) into frequencies of different cell types, including stem cells, from bulk RNA-Seq data. b. Cancer stem cell identification workflow using a machine learning-based identifier from an input of labeled genomic signatures of different stem cell types. c. Comparative performance analysis of different ML models used to identify stem cells. The dataset was randomly divided into training and test sets for five different runs in a 70:30 ratio. Five different performance parameters were used, as indicated in the figure. d. Representative single-run confusion matrix of true level vs. predicted level showing the prediction accuracy of SVM used for cancer stem cell identification. In each run, the dataset was randomly divided into training and test sets at a 70:30 ratio. e. Identifier model performance was assessed by measuring the area under the curve (AUC) of the ROC curve and quantifying the true-positive and false-positive rates for SVM. The ROC-AUC values for each CSC state are mentioned within the figure.

Utilizing provided RNAseq data samples, the OSCORP identification module predicts distinct cancer stem cell (CSC) states within tumors. These states, induced-pluripotent (iPCSC), embryonic stem cell-like (ECSC), multipotent (MCSC), and unipotent (UCSC), are classified based on their genetic similarity to normal human stem cell states (iPSC, hESC, hMSC, and hUSC, respectively) (Fig. 1b). The identifier module was trained using the Stemformatics dataset[15], encompassing labeled mRNA profiles of different stem cell states (Extended Data Fig. 1c, n = 3294, power = 1). Although iPSCs and hESCs share pluripotency, their distinct global genomic signatures[34] (Extended Data Fig. 1d-1 g) are maintained as separate classes to preserve their intrinsic genomic characteristics, aligning with CSC origin hypotheses[35,36]. In this context, iPCSC signifies CSCs originating from cancer seed cells through induced transcriptomic rewiring, while ECSCs represent tissue (adult) stem cells acquiring mutations to become pluripotent CSCs. Six machine learning models—logistic regression (LR), support vector machine (SVM), decision tree (DT), random forest (RF), multilayer neural network (NN), and XGBoost (XGB)—were employed for training (Fig. 1c, Extended Data Fig. 2). Rigorous k-fold cross-validation (k=5) validated SVM as the top-performing identifier model (Fig. 1c, 1d; Extended Data Fig. 2a). OSCORP achieved a weighted F1 score of 0.98 across all stem cell types, indicative of well-balanced precision and recall of 0.98 (Extended Data Fig. 2b). NN and LR also demonstrated weighted F1, precision, and recall scores of 0.98. Despite acceptable ROC-AUC values for all models except RF (Fig. 1e, Extended Data Fig. 2c), SVM was chosen for its consistent prediction performance across multiple runs (Fig. 1c). OSCORP maintains a consistent accuracy of approximately 99% (weighted accuracy = 98.67%) across various CSC states, demonstrating robustness and reliability, even in the face of inherent class imbalance within the dataset.

**Figure 2.**
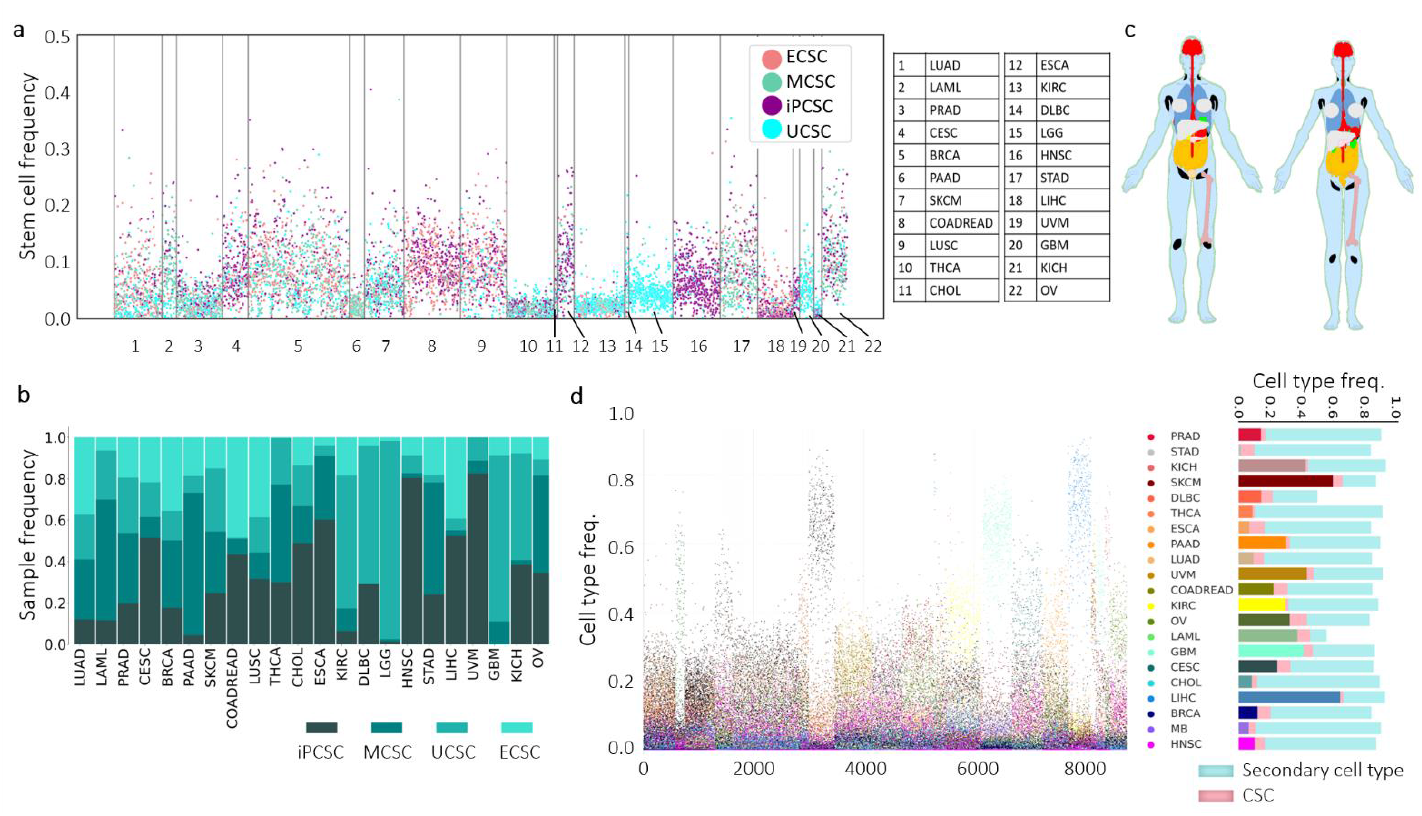
Panorama of pancancer stem cell distribution and CSC-induced heterogeneity. a. Pancancer panorama of stem cell state frequencies. b. The distribution of stem cell subtypes within various solid tumor categories shown in terms of the frequency of patients affected. c. Anatograms demonstrating the organ-specific dispersion of CSC subtypes within solid tumors, independently for males and females. d. The cancer-specific allocation of diverse cell-type frequencies within a given sample as a distribution of different cell types depicting the intratumor heterogeneity induced by cellular plasticity. The corresponding vertical stacked bar plot displays the cancer-specific frequencies of distinct cell types residing in tumors. This includes primary tumor cells, secondary cell types, and stem cell frequencies across different solid tumors. Infiltrating immune cells are not considered here.

### Pancancer profiling of CSCs using OSCORP

Our examination of 22 solid tumor types within the TCGA dataset unveiled a distribution of stem cell frequencies ranging from 1% to 25% (Fig. 2a), aligning with findings from diverse experiments utilizing different assay types[37,38]. Notably, none of the patients in our analysis lacked CSCs, underscoring the pervasive role of CSCs in the trajectory of cancer progression (Fig. 2a). Intriguingly, both GBM and LGG samples exhibited an absence of the iPCSC state. One conceivable explanation for this absence may lie in the distinctive characteristics of primary brain tumors, attributed to the unique cellular environment and the presence of the blood‒brain barrier, which collectively may hinder the development of iPCSCs. Furthermore, the observation could be influenced by sample size and patient selection, as the limited sample size might have insufficiently captured their presence.

Exploring the distribution of stem cell states across sexes, we observed no noteworthy distinctions between males and females (Fig. 2b, Extended Data Fig. 3a). Nevertheless, females exhibited elevated stem cell frequencies across diverse states (Extended Data Fig. 3b). This disparity may stem from hormonal variations, where estrogen could influence stem cell activity, coupled with distinctions in the tumor microenvironment and immune responses between the sexes. Additionally, we observed a surge in stem cell frequency with advancing age (Extended Data Fig. 3c), likely attributed to age-related alterations in the tissue microenvironment and immune system. Such changes may foster CSC expansion, heightening susceptibility to age-related cancers[39]. Notably, CSC frequencies exhibited positive correlations with the MSI-MANTIS score and aneuploidy score, serving as indicators of genomic instability (Extended Data Fig. 3d, 3e). The aneuploidy score displayed a decreasing pattern from ECSC to UCSC (Extended Data Fig. 3f), underscoring its significance in CSC differentiation. This intricate interplay of stem cell dynamics across sexes, ages, and genomic instability indicators contributes to a nuanced understanding of the underlying factors influencing CSC behavior in diverse contexts.

**Figure 3.**
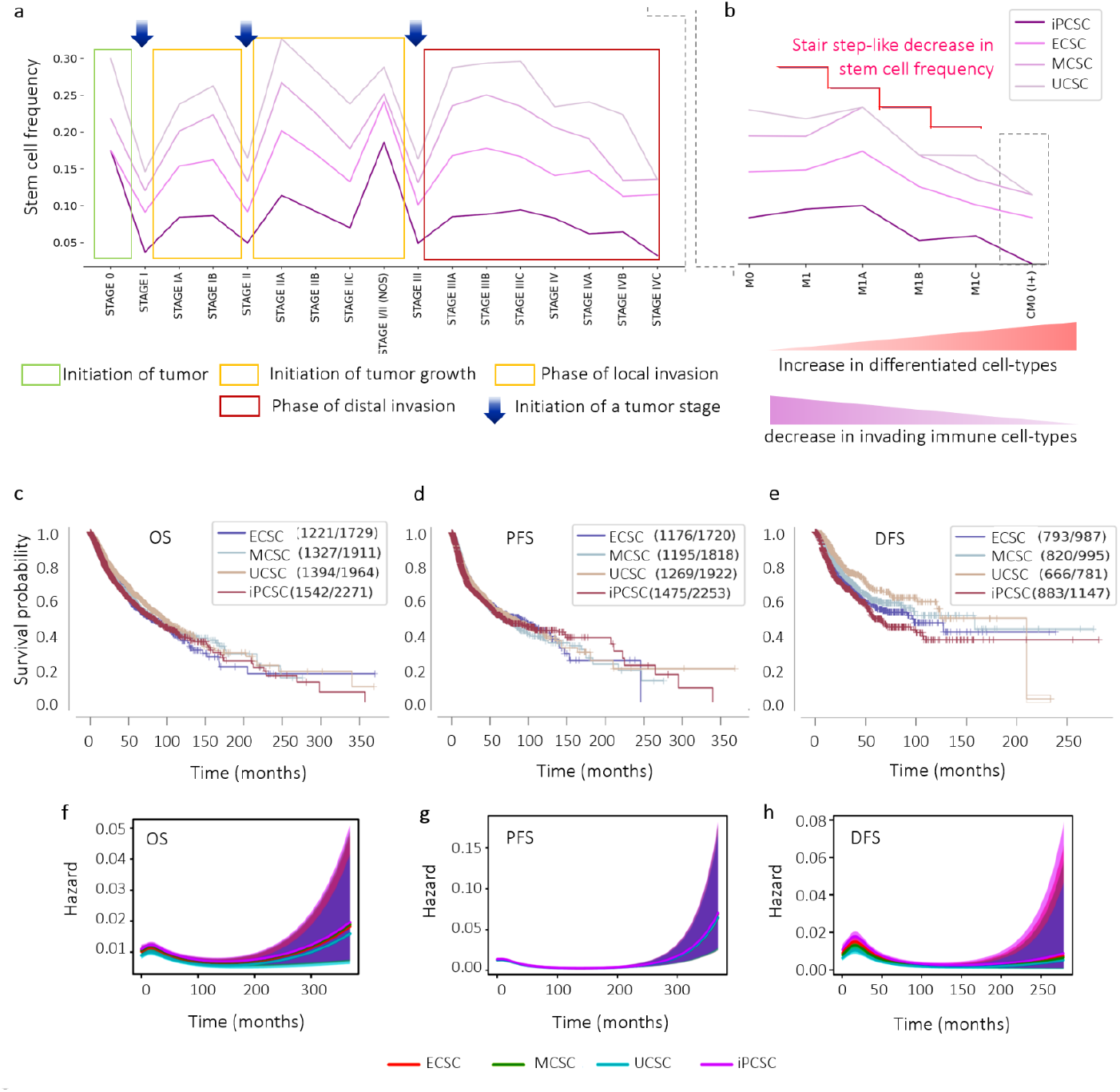
Importance of CSC states in cancer clinical events and survivability. a. Distribution of frequencies of the different CSC states along the AJCC tumor stages in the pancancer cohort. Initiation of each stage is indicated with a downward arrow, whereas rectangular blocks mark the progression of each stage denoting major cancer events. b. Distribution of frequencies of the different CSC states along the metastatic stages in the pancancer cohort. Changes in differentiated cell type frequency and infiltrating immune cell frequencies are shown below. c. KM plot showing how different CSC states affect the overall survival of cancer patients. d. The effect of CSC states associated with patient tumors on progression-free survivability. e. The effect of CSC states associated with patient tumors on disease-free survivability, which also indicates tumor relapse. (f-h) Continuous hazard function prediction for overall, progression-free, and disease-free survivability using the Cox regression model.

We explored the distribution of diverse stem cell states across different tumor categories, assessing patient prevalence (Fig. 2c), thereby revealing the intricate landscape of tumor heterogeneity arising from the differentiation of cancer stem cells (CSCs). This phenomenon gives rise to multiple subpopulations within tumors (Fig. 2d). The coexisting subpopulations predominantly exhibited negative correlations, with their dynamics showcasing uniqueness specific to each cancer type (Extended Data Fig. 4a). Notably, genetically similar cancer types demonstrated positive correlations in terms of coexistence (Extended Data Fig. 4b). Utilizing a Gaussian graphical model to visualize the intricate relationships between primary and secondary tumor types (Extended Data Fig. 4c), we observed that PRAD, ESCA, CHOL, and BRCA displayed a prominent degree of coexistence with various secondary tumor types, potentially indicating heightened heterogeneity within these cancer types. This comprehensive exploration provides valuable insights into the complex dynamics of CSC differentiation and its impact on tumor heterogeneity across a spectrum of cancer categories.

**Figure 4.**
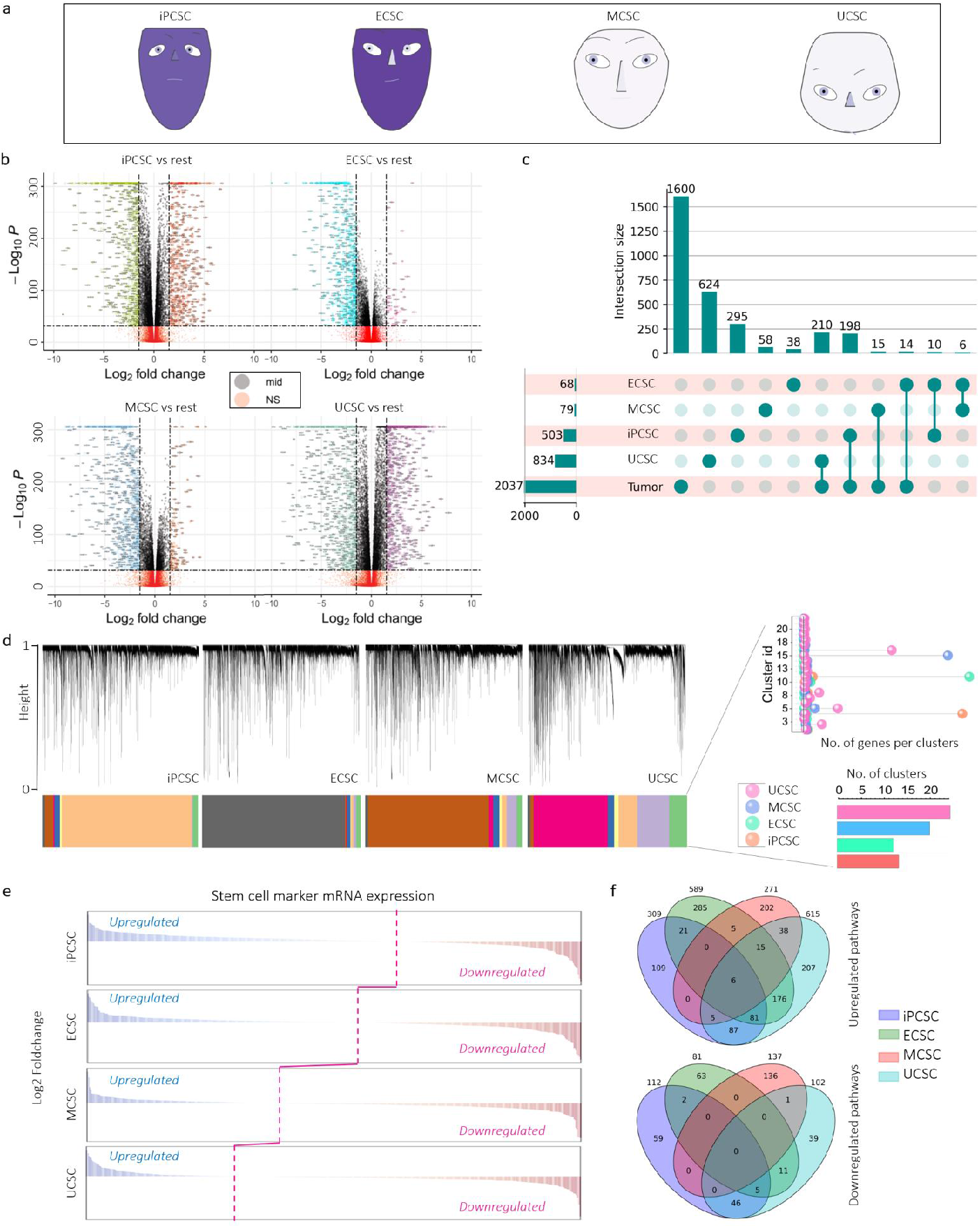
Differential gene expression patterns define distinct CSC states. a. Chernoff’s face for different CSC classes showing distinct global genomic signatures within those particular classes. Facial properties such as face shape, face color, the shape of eyes, eyebrows, lips, and nose are indicative of mRNA expression patterns of gene clusters. Gene expression values were taken as log2(CPM+1). b. Volcano plots showing the significantly (p-val < 10e-32) over/underexpressed genes in each CSC class concerning their log2-fold change (significance level > 1.5). c. Overlap of overexpressed genes in different CSC states. The uniquely overexpressed genes can contribute as markers for corresponding states. WGCNA of gene clusters obtained from gene regulatory networks for each CSC type. The clustering dendrogram and expression heatmap of genes identifying the WGCNA modules are shown. Gene clustering was based on TOM-based dissimilarities. The corresponding lollipop plot shows the number of genes per cluster in each stem cell type, while the bar plot compares the number of clusters in CSC classes. e. Gene Set Enrichment Analysis showing RNAseq-based upregulated and downregulated gene sets representative of cancer stemness. The pink line passes through the origin of each curve and denotes the sequential change in the expression pattern of stemness markers from pluripotency to unipotency. f. Number of uniquely and commonly upregulated and downregulated pathways in different CSC states.

### Analyzing the impact of CSCs on cancer-associated clinical characteristics

We conducted an in depth, CSC-specific examination of TCGA patient data (n = 8786). Intriguingly, we observed a cyclical pattern in CSC frequency at the onset of each tumor stage, showing an initial drop followed by a steady increase during that stage before eventual decline (Fig. 3a). This cyclic behavior suggests periodic depletion and replenishment of CSC subpopulations as cancer progresses. Stem cell frequency gradually decreases during metastatic phases, accompanied by an increase in differentiated cell types and a decrease in infiltrating immune cells (Fig. 3b), indicating a potential shift toward more specialized cancer cells contributing to metastasis [4,19]. The rise in differentiated cell types and reduced immune cell frequencies correspond to the evolving tumor microenvironment, possibly impeding immune infiltration and promoting cancer progression [40,41]. Tumor grade-specific analysis revealed an increase in stem cell frequency from grade 1 to 3, followed by a sharp decrease in grade 4 (Extended Data Fig. 5a), suggesting a transition from undifferentiated CSCs to more aggressive, differentiated CSCs during cancer progression.

**Figure 5.**
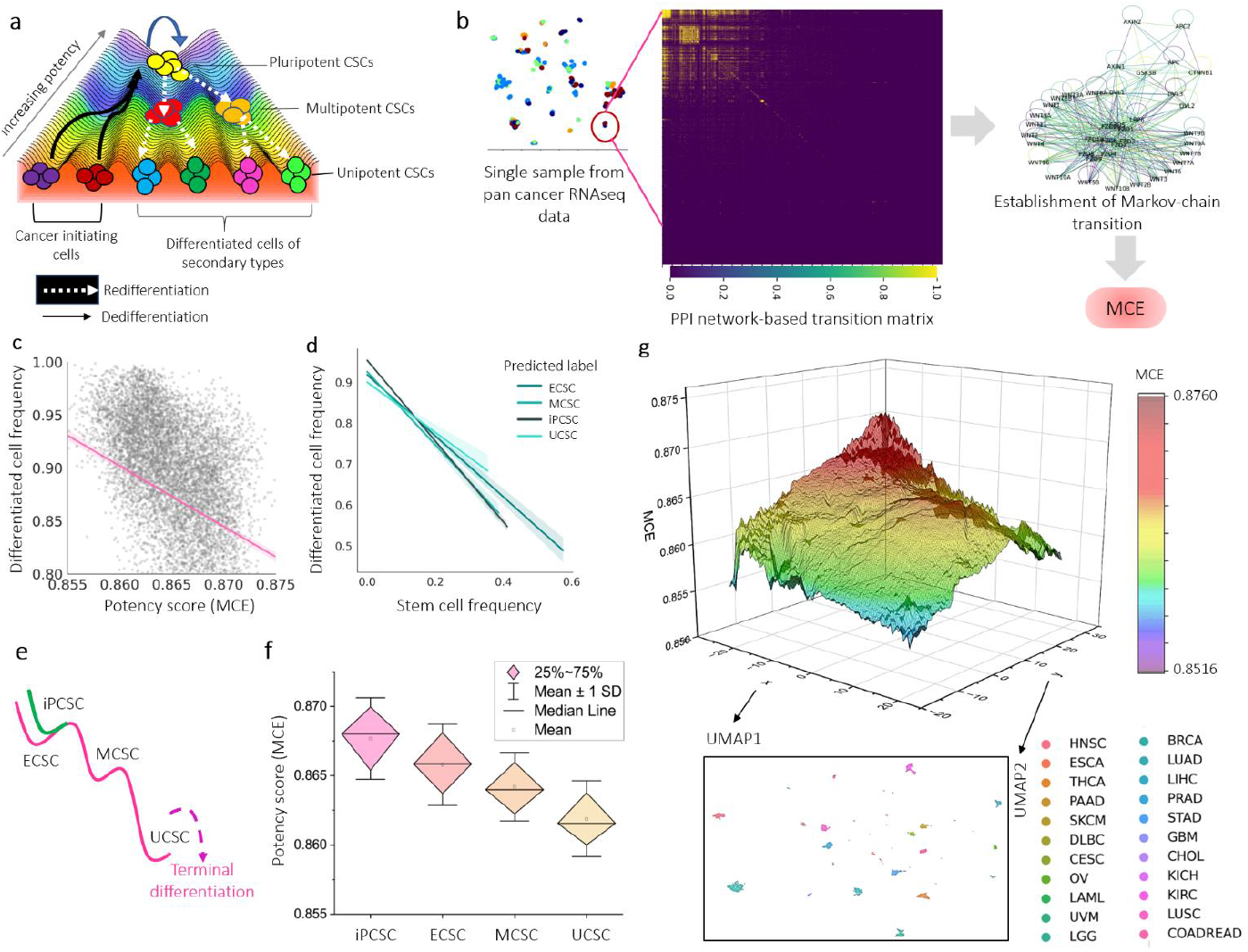
Markov chain entropy-based method to shape the pancancer potency landscape of tumor-associated stem cells. a. A simplistic model of CSC dedifferentiation from cancer-initiating cells and their gradual differentiation to CSC states with lower potency over a Waddington-like epigenetic landscape. From the bottom to the top of the hill, cellular potency increases. b. Workflow to determine each tumor sample’s Markov chain entropy (MCE) as a quantitative indicator of cellular potency from the PPI network-based transition matrix developed using the corresponding gene expression signature. c. Correlation between the frequency of differentiated cell types and the potency score (MCE) of tumor samples. correlation between the frequency of differentiated cell types and the state-specific CSC frequency for tumor samples. e. Hypothetical side view of a classical Waddington-like epigenetic landscape showing the positioning of CSC states as obtained from our identifier model. f. For RNA-Seq dataset profiling over 8000 cancer samples across 22 solid tumor types classified into four different CSC states, we show corresponding box plots of the predicted potency scores for MCE as a potency measure. g. A pancancer Waddington-like epigenetic landscape composed of ~8000 tumor samples across 22 solid tumor types. The curve was simulated using the nonlinear logistic function (maximum likelihood) to increase the points 100 times. The Z-axis corresponds to the MCE values, whereas the X- and Y-axes show the two dimensions of UMAP obtained from supervised dimensionality reduction of the RNAseq data of the samples according to their cancer type. The UMAP is shown separately.

Examining the connection between various CSC frequencies and posttreatment tumor recurrence (Extended Data Fig. 5b), we found that differentiated CSCs (MCSCs and UCSCs) were significantly more common in tumor recurrence cases than in nonrecurrent patients (p-val=0.004 and 0.016, respectively). In contrast, the frequencies of pluripotent CSCs (ECSC and iPCSC) remained relatively stable (p-val=0.634 and 0.952, respectively). Pluripotent CSCs in recurrent tumors may conceivably differentiate into specialized cell types to meet evolving tumor requirements, augmenting the frequency of differentiated CSCs. As shown in Extended Data Fig. 5c, the presence of CSCs significantly affected the chemotherapeutic response (p-val=0.0018), with a notable increase in CSCs, except iPCSCs (p-val=0.731), in patients reporting a better clinical response. The response to radiation therapy showed a similar pattern (Extended Data Fig. 5d), except for lowered radiation effectiveness against ECSCs (p-val=0.025). This impact of CSCs on clinical response and recurrence is evident in CSC-specific survival outcome analyses. Although iPCSCs had the highest effect on overall survival (Cox p-val=0.0003) (Fig. 3c), progression-free survival showed no CSC-specific effects (Fig. 3d). Disease-free survival (DFS), an indicator of tumor recurrence, is notably affected by CSC states, with iPCSCs and ECSCs showing poorer survival outcomes (Cox p-val=1.5×10^−6^ and 0.0001, respectively) (Fig. 3e) and serving as prognostic indicators of cancer relapse. A similar trend was evident in the temporal survival hazard analysis of different CSC states (Fig. 3f, g, h), where pluripotent stem cells exhibited high hazard values for overall survival and disease-free survival, underscoring their significance in cancer prognosis.

### Global genomic signatures of CSC states unveiled by OSCORP

Examining the distinctive genomic profiles within the hierarchy of tumor-resident stem cells (CSCs), we extended our investigation beyond clinical traits to unravel the genetic intricacies governing these characteristics. Our in-depth gene expression analysis unveiled a complex genomic diversity among predicted CSC states, depicted through the Chernoff face representation (Fig. 4a). Notably, tumor-resident induced pluripotent stem cells (iPCSCs) and tumor-resident epithelial stem cells (ECSCs) share a similar global genomic signature, setting them apart from tumor-resident mesenchymal stem cells (MCSCs) and tumor-resident unipotent stem cells (UCSCs). iPCSCs exhibited a significant number of overexpressed and downregulated genes compared to other states, while ECSCs displayed a relatively low number of overexpressed genes, akin to UCSCs (Fig. 4b). In contrast, MCSCs demonstrated substantial up- and downregulation of numerous genes compared to the rest. Unique overexpressed genes were identified in each of these four states, with 38, 58, 295, and 624 genes specific to ECSCs, MCSCs, iPCSCs, and UCSCs, respectively (Fig. 4c, p-val<10^−32^). Notably, these genes are exclusive to their respective CSC states, eliminating interference from malignant cell-specific expression. Interestingly, shared genes between iPCSCs and malignant cells were identified, potentially indicating their origin from dedifferentiated malignant cells.

Utilizing weighted gene coexpression network analysis (WGCNA), we identified highly correlated genes among meta-genes in the four CSC states. The gene expression correlation matrix was transformed into a scale-free topology network, revealing modules of hub gene networks represented by high mean connectivity in each CSC state (Extended Data Fig. 6a). WGCNA highlighted highly correlated gene modules in ECSCs (10 and 12 modules), iPCSCs (17 and 21 modules), MCSCs, and UCSCs (Fig. 4d). Hierarchical clustering and dynamic tree-cut analysis revealed significant module shifts from ECSC to UCSC, indicating a progressive alteration, while iPCSCs displayed a distinct clustering pattern. The smaller information entropy in pluripotent CSCs, evidenced by fewer clusters, implies less diverse and more predictable gene expression patterns, whereas MCSCs and UCSCs displayed increased gene cluster diversity, reflecting a complex and heterogeneous gene expression landscape.

**Figure 6.**
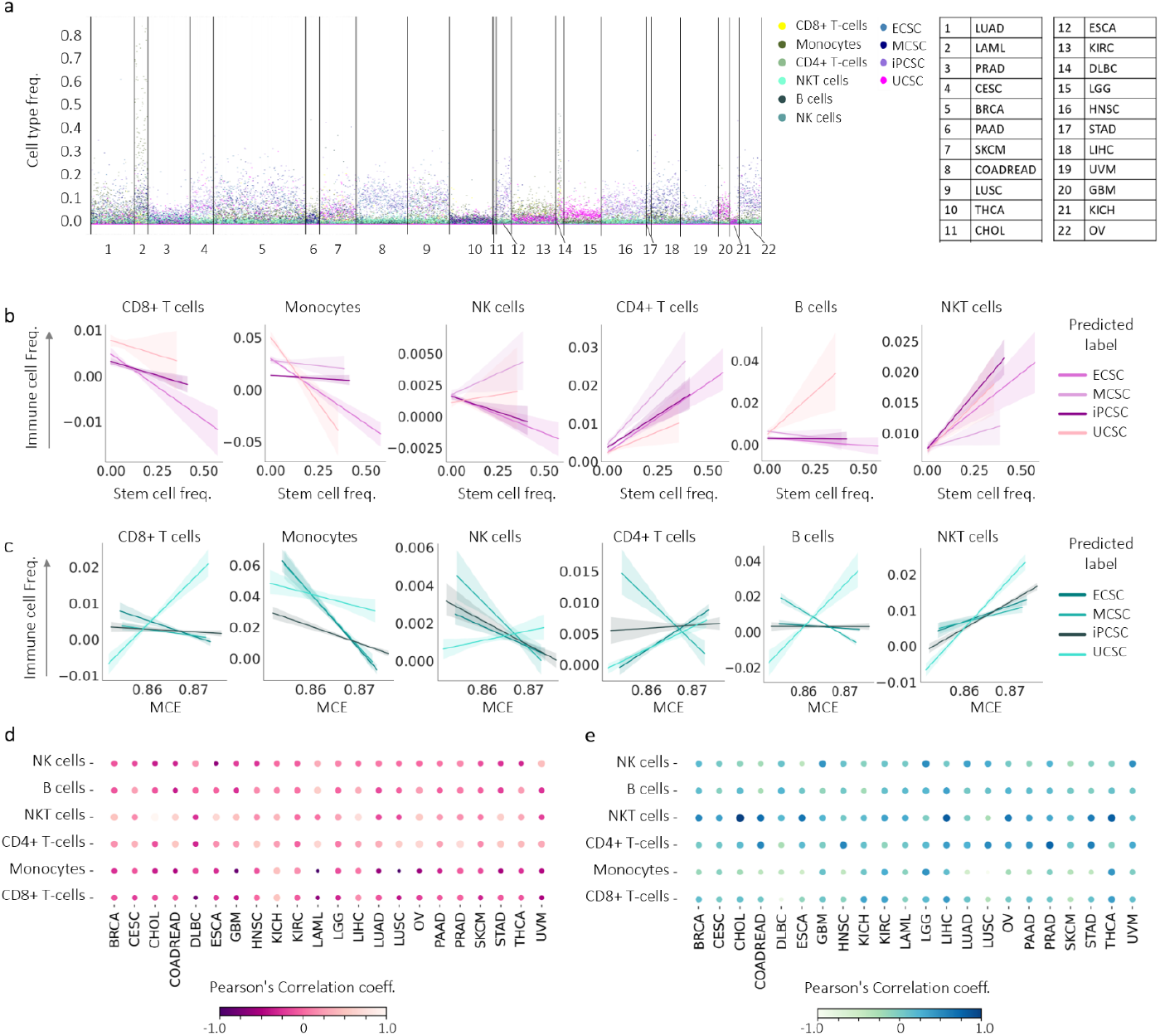
The presence of CSCs affects the landscape of tumor-infiltrating immune cells. a. Panorama of tumor-associated stem cell abundance of corresponding states and associated immune infiltration frequencies for 22 solid tumor types. Colors are indicative of different cell types as mentioned in the legend. Each column represents a particular tumor type. b. Correlation of tumor-infiltrating immune cell frequencies with stem cell frequencies in different CSC subgroups. c. Correlation of tumor-infiltrating immune cell frequencies with potency score (MCE) in different CSC subgroups. d. Cancerwise bubble plot showing Pearson’s correlation coefficient of tumor-infiltrating immune cell frequencies with stem cell frequencies. e. Cancerwise bubble plot showing Pearson’s correlation coefficient of tumor-infiltrating immune cell frequencies with MCE.

Our differential gene expression analysis of validated CSC markers corroborates the transition pattern from pluripotent to unipotent CSCs (Fig. 4e). The gradual decrease in the overexpression of CSC markers [14,28] from iPCSC to UCSC confirms the progressive differentiation in CSCs, mirroring normal stem cell behavior. Using WGCNA modules of eigengenes that exhibit similar expression patterns across samples, we studied CSC state-specific gene clustering (Extended Data Fig. 6c). This grouping highlights coregulated genes that might be involved in similar biological processes. Pathway enrichment analysis of these modules highlighted pluripotency-promoting pathways, such as Wnt, Hippo, Notch, Hedgehog, and HIF-1α signaling in pluripotent CSCs, which are downregulated in MCSCs and UCSCs (Extended Data Fig. 7a-d). We identified 109, 285, 202, and 207 differentially upregulated and 59, 63, 39, and 46 downregulated pathways for iPCSC, ECSC, MCSC, and UCSC, respectively (Fig. 4f, p-val<0.05, FDR q-val<0.05). Enrichment analysis with Gene Ontology (GO) biological processes indicated a gradual enrichment of cellular differentiation-specific processes and related genes from iPCSCs/ECSCs to UCSCs (Extended Data Fig. 7e-7 h, p-val<0.05, FDR q-val<0.05). The enrichment of purine and amino acid metabolism appears to favor differentiation, whereas pluripotency is enriched in G protein-coupled receptor (GPCR) signaling and cholesterol metabolism. The pyruvate dehydrogenase complex and tricarboxylic acid (TCA) cycle are downregulated with progressive potency loss. This intricate interplay between CSC plasticity and signaling alterations underscores the dynamic nature of different CSC states.

### OSCORP reveals a pancancer potency landscape of CSC differentiation/dedifferentiation

The intrinsic genomic signatures and their interplay with the tumor microenvironment intricately govern the potency of tumor-resident stem cell (CSC) states [44], defining their capacity to differentiate into multiple lineages, as illustrated in the stem cell potency landscape (Extended Data Fig. 8a). This landscape serves as a topographic map shaped by gene expression patterns guiding cells toward specific fates (Fig. 5a). In our investigation, we employed Markov chain entropy (MCE) to quantify CSC potency, assuming that gene expression arises from an unknown Markovian signaling process estimated through a network model of signaling interactions [45] (Fig. 5b). The sorting of samples by MCE as a proxy for CSC potency revealed a negative correlation with differentiated stem cell frequency (Fig. 5c, r=-0.247, p-val=6.172×10^−121^). Consistent with expectations, induced pluripotent stem cells (iPCSCs) and embryonic stem cells (ECSCs) exhibited a stronger negative correlation, while unipotent stem cells (UCSCs) displayed the lowest (Fig. 5d). Following the consensus in stem cell biology, the order of MCE was found to be iPCSCs > ECSCs > multipotent cancer stem cells (MCSCs) > UCSCs, outlining the trajectory of CSC differentiation (Fig. 5e, f). The remarkable alignment between our identification of CSC states and their potency score underscores the robustness of our CSC identification model.

A reduced MCE, indicating more differentiated tumor cells and contributing to diverse malignant subpopulations, enabled the construction of a pancancer potency landscape of CSCs. Inspired by Waddington’s concept, this landscape combines MCE on the z-axis with cancer sample positions on the x- and y-axes using UMAP-derived reduced dimensions (Fig. 5g). High-potency states such as iPCSCs and ECSCs are positioned at higher altitudes, while valleys represent less differentiated, specialized CSCs such as UCSCs. This landscape effectively captures the transition of CSCs from an undifferentiated to a lineage-specific state, illustrating their journey toward specification. A parallel analysis using another potency metric, stem cell entropy (SCENT) [46], yielded consistent results, affirming the model’s robustness (Extended Data Fig. 8b, c). The strong positive correlation between MCE and SCENT (r=0.941, p-val=0) further validates the accuracy of our model (Extended Data Fig. 8d).

### Effect of CSCs on tumor immune infiltrates

CSCs possess the capability to evade immune detection and foster an anti-inflammatory, protumorigenic immune environment, thereby facilitating cancer progression [47]. Notably, tumor-associated macrophages contribute to the growth of CSCs through STAT-mediated positive feedback [47]. However, the outcomes of interactions between immune cells and CSCs exhibit a mixed nature, with some interactions promoting CSCs while others hindering them [47]. To delve into the dynamics of these CSC-immune cell interactions, an investigation into the coenrichment of immune cells with various CSC states across 22 cancer types was conducted (Fig. 6a). The findings revealed that CD4+ T cells and NKT cells coenriched with most CSC states, especially in pluripotent CSC samples, where higher immune cell frequencies were observed, particularly in monocytes, NK cells, and B cells. Positive correlations were noted between CD4+ T cells (r=0.18; p-val=6.85×10^−44^) and NKT cells (r=0.13; p-val=1.95 × 10^−30^) with various CSC states, while negative correlations were observed for CD8+ T cells (r=0.86; p-val=5.67×10^−16^) and monocytes (Fig. 6b). Interestingly, the comparative analysis of immune cell frequencies concerning the sample-specific potency score (MCE) revealed varying trends, such as decreasing monocyte (r=-0.201; p-val=6.867×10^−81^) and NK cell (r=-0.113; p-val=1.038×10^−8^) frequencies with increasing MCE in pluripotent CSCs, while NKT cell frequencies increased significantly (r=0.243; p-val=5.798×10^−34^) (Fig. 6c). Notably, the overexpression of known CSC markers (mRNAs) did not consistently correlate with the relationship between CSCs and immune cells, suggesting intricate cancer-specific variations [Extended Data Fig. 10]. In conclusion, the intricate interactions between CSCs and the immune microenvironment have profound implications for cancer progression and targeted therapeutic interventions, revealing a complex network of immune cells influencing CSC states in various cancer types.

## Discussion

The exploration of tumor-resident stem cells (CSCs) has garnered significant attention in recent years, driven by their pivotal role in cancer progression and therapeutic resistance [1,5,6,8,17,19,21,26,41]. While traditional methods have contributed to our understanding of CSC biology, inherent limitations underscore the significance of OSCORP. Biomarker-based assays, commonly employed for CSC identification, face challenges from tumor-microenvironment interactions and regulatory factors, introducing avoidable interference [9,28,48]. Colony-forming assays, immunohistochemistry, density gradient centrifugation, and FACS methods exhibit variability in CSC identification, causing disparities across runs or operators [9,12,49,50]. Xenograft assays, due to complex engraftment procedures and varying protocols, yield inconsistent results, emphasizing OSCORP’s advantage in minimizing technical variability through single-source RNAseq data per case. OSCORP, leveraging computational cytometry, offers high-throughput, simultaneous CSC profiling across multiple cancer types, addressing scalability and resource-intensive limitations associated with traditional wet-lab methods.

A transformative aspect of OSCORP lies in its ability to unveil the diversity among CSCs. Past studies often treated CSCs as a relatively uniform potency state, overlooking intricate variations within [13,51]. In contrast, OSCORP classifies CSCs into four distinct states, allowing extensive exploration of molecular signatures linked to each CSC state. Capturing the dynamic behavior of CSCs, a challenge for conventional techniques is addressed by OSCORP through Markov chain entropy (MCE) as a measure of cellular potency. This approach provides insights into how CSCs navigate a complex cellular landscape, offering a detailed map of their journey through various potency states. OSCORP delivers a comprehensive view of CSC diversity and its consequences for clinical outcomes, revealing previously unnoticed elements such as the sex-specific distribution of CSC states and their relevance in disease-free progression.

While OSCORP signifies a substantial advance in CSC research, the need for large-scale time-lapse RNAseq data is underscored. The synergy between such data and computational approaches, as demonstrated by OSCORP, holds promise for more comprehensive insights into CSC biology and clinical implications. It is essential to acknowledge that our model cannot distinguish between CSCs and tumor-resident normal stem cells or account for potential interconvertibility between iPSCs and ECSCs due to the lack of suitable scRNAseq data for model training [52]. However, this is not an inherent problem with our model, and OSCORP can address this with relevant data access.

Considering these factors, OSCORP emerges as a potential solution to the shortcomings of conventional laboratory techniques in CSC quantification, exhibiting near-absolute accuracy. Its high-throughput analysis of CSC dynamics by categorizing them into distinct yet dynamic states positions it as a transformative tool for the field. We envision OSCORP as a potential clinical tool for monitoring patient-specific CSC changes during treatment when paired with routine sequencing data. With future advancements, this AI-driven approach holds the potential to unravel the complexities of CSC biology, ushering in a more precise and efficient paradigm for cancer research and treatment.

## Data Availability

All datasets used in this study are publicly available. The raw count TCGA data can be found at the Broad Institute FireBrowse portal (https://gdac.broadinstitute.org/). The raw stem cell RNA-seq data can be obtained from Stemformatics (https://www.stemformatics.org/). The raw stem cell scRNA-seq data can be obtained from GEO with accession IDs GSE66507 and GSM3370006. The tumor scRNA-seq data can be obtained from TISCH2 (http://tisch.comp-genomics.org/). The immune cell single-cell RNA-Seq data can be obtained from the PBMC Zenodo page (https://zenodo.org/records/4730807). All intermediate data supporting the findings of this study are available from the corresponding authors upon reasonable request.

## Code Availability

All codes used in this study are available from the corresponding author upon reasonable request.

## Acknowledgments

The results published here are based upon data generated by the TCGA, STEMFORMATICS, and TISCH2 projects, and we are grateful to these consortia for their contributions. We thank S.N. Bose National Centre for Basic Sciences and Ashoka University for support and funding. S.H. thanks DST SERB Core Research Grant for funding. D.C. thanks DBT for funding and support through the DBT-JRF fellowship. B. N and D. G are thankful to the Mphasis Foundation for their support. The funders had no role in the study design, data collection and analysis, decision to publish or preparation of the manuscript.

## Author Contributions

S.H. and D.G. conceived the study. D. C, B. N, S. H, and D. G designed the study. S. P, and S. M curated the data. S. P and D. C developed the quantifier model. B. N and S. M developed the identifier model. S. P and D. C designed the potency score module. D. C, D. M, and S. P performed the data analysis. D. C, D. M, S. P, B. N and S. M wrote the manuscript. All the authors took part in interpreting the findings and reviewing the manuscript.

